# Mitotic chromosomes harbor cell type and species-specific structural features within a universal looping architecture

**DOI:** 10.1101/2023.12.08.570796

**Authors:** Marlies E. Oomen, A Nicole Fox, Inma Gonzalez, Amandine Molliex, Thaleia Papadopoulou, Pablo Navarro, Job Dekker

## Abstract

The architecture of mammalian mitotic chromosomes is considered to be universal across species and cell types. However, some studies suggest that features of mitotic chromosomes might be cell type or species specific. We previously reported that CTCF binding in human differentiated cell lines is lost in mitosis, whereas mouse embryonic stem cells (mESC) display prominent binding at a subset of CTCF sites in mitosis. Here, we perform parallel footprint ATAC-seq data analyses of mESCs and somatic mouse and human cells to further explore these differences. We then investigate roles of mitotically bound (bookmarked) CTCF in prometaphase chromosome organization by Hi-C. We do not find any remaining interphase structures such as TADs or CTCF loops at mitotically bookmarked CTCF sites in mESCs. This suggests that mitotic loop extruders condensin I and II are not blocked by bound CTCF, and thus that any remaining CTCF binding does not alter mitotic chromosome folding. Lastly, we compare mitotic Hi-C data generated in this study in mouse with publicly available data from human and chicken cell lines. We do not find any cell type specific differences; however, we do find a difference between species. The average genomic size of mitotic loops is much smaller in chicken (200-350 kb), compared to human (500-750 kb) and mouse (1-2 mb). Interestingly, we find that this difference in loop size is correlated with the average genomic length of the q-arm in these species, a finding we confirm by microscopy measurements of chromosome compaction. This suggests that the dimensions of mitotic chromosomes can be modulated through control of sizes of loops generated by condensins to facilitate species-appropriate shortening of chromosome arms.

## Introduction

The development of 3C-techniques (Dekker et al. 2002; Dostie et al. 2006; Lieberman-Aiden et al. 2009; Belaghzal et al. 2017) has contributed to a better understanding of key features of chromosome organization in vertebrate cells. Interphase chromosomes are organized on the megabase scale in A and B compartments and on a smaller scale of tens to hundreds of kilobase in topologically associating domains (TADs) (Lieberman-Aiden et al. 2009; Erdel and Rippe 2018; Michieletto et al. 2016; Dixon et al. 2012; Rao et al. 2014; Nuebler et al. 2018; Nora et al. 2012). TADs are proposed to be formed by loop extruding machines, such as cohesins, which can be blocked by the chromatin binding protein CCCTC-binding factor (CTCF) when bound to its motif (Fudenberg et al. 2016; Nora et al. 2016; Rao et al. 2017, 2014; Dekker and Mirny 2016; Nuebler et al. 2018; Sanborn et al. 2015; Wit et al. 2015). Although the mechanisms that establish and maintain these structures are largely shared between different cell types and between different vertebrate species, the specific genomic regions that interact can differ strongly between species, cell types, and even between sick and healthy cells (Oksuz et al. 2020; Smith et al. 2016; Lupianez et al. 2015; Valton and Dekker 2016; Rao et al. 2014; Dekker and Mirny 2016).

In contrast to interphase chromatin, vertebrate mitotic chromosomes are often perceived as universal structures, independent of cell type or organism. Historically studied by microscopy (Earnshaw and Laemmli 1983; Marsden and Laemmli 1979; Flemming 1878) and in more recent years using genomics techniques (Gibcus et al. 2018; Naumova et al. 2013; Abramo et al. 2019), we have gained understanding on the fundamental principles of mitotic chromosome folding. In mitosis, the interphase structures are completely dissolved, as both TADs and compartments can no longer be observed (Naumova et al. 2013; Gibcus et al. 2018). Instead, chromosomes are folded as helical loop arrays mediated by condensin I and II, which are not positioned at any specific genomic locations (Belmont 2006; Batty and Gerlich 2019; Gibcus et al. 2018). This results in the observation of a generally smooth inverse relationship between genomic distance and interaction frequency without any site-specific features, when studying mitotic chromosomes in cell populations by Hi-C (Gibcus et al. 2018; Naumova et al. 2013).

This might give the impression that mitotic chromosomes in all biological contexts are organized in a similar fashion. However, microscopy and biochemical studies revealed that condensins play a more complex role during the rapid cell cycle of mouse embryonic stem cells (mESCs) (Fazzio and Panning 2010). It has been shown in *Xenopus leavis* that mitotic chromosomes from sperm nuclei are folded as long and thin structures but become increasingly shorter and fatter throughout the early stages of development (Kieserman and Heald 2011). Additionally, depletion experiments in *Xenopus leavis* extract experiments show that the ratio of condensin I and II can affect the width-to-length ratio of chromosomes in mitosis (Shintomi and Hirano 2011; Zhou et al. 2023). Along these lines, it has been described recently that the degree of chromosome arm compaction during mitosis can differ across species (Kakui et al. 2022).

Using genomics techniques, it was found that mitotic chromosomes can harbor cell type-specific features on a more detailed scale, e.g., in chromatin accessibility at the level of the nucleosomal array, histone modifications, and mitotically bound chromatin factors (Oomen et al. 2019; Festuccia et al. 2016; Wang and Higgins 2013; Hsiung et al. 2015). Of particular interest are studies that found that the architectural protein CTCF remains bound to a subset of its binding sites during mitosis in some cell lines, while it is completely displaced in others: In differentiated human cell lines HeLa and HFF, we have previously reported complete loss of CTCF binding by ATAC-seq, Cut&Run and imaging (Oomen et al. 2019). Similarly, we described complete loss of binding in the mouse somatic cell lines C2C12 and 3T3 (Owens et al. 2019). In contrast, we showed in mESC that a substantial fraction of CTCF sites remains bound in mitosis (Owens et al. 2019), and this persistent association has been linked to CTCF-dependent post-mitotic reactivation of a small subset of promoter-restricted mitotic CTCF targets (Chervova et al. 2022). Moreover, mitotic CTCF binding was also associated with faster reassembly of 3D contacts during early interphase of pluripotent cells (Pelham-Webb et al. 2021). These observations are in line with independent observations in a mouse blood progenitor cell line, in which the retained CTCF binding has been implicated in faster transcription reactivation, when involving promoters, and more generally in fast restoration of 3D contacts after mitosis (Zhang et al. 2019). Together, these reports suggest that mitotic chromosomes are not strict universal structures across eukaryotes, and that the overall dimensions of the mitotic loop array arrangement as well as the local chromatin state can reflect both species-specific features as well as characteristics of its cell type identity.

In this study, we first performed parallel footprinting analyses of ATAC-seq data to confirm that mitotic CTCF binding is prominent in mESCs only. Notably, comparative Hi-C analyses did not unmask any conformational specificity associated to mitotic CTCF binding, indicating that mitotically retained CTCF sites do not influence condensin-mediated loop extrusion and mitotic chromosome formation. Interestingly, these analyses revealed species-specific differences in mitotic chromatin loop sizes in relation to differences in genomic arm length. We find that mitotic chromosome architecture is insensitive to species and cell type-dependent differences in CTCF retention, and is adaptable through modulation of loop sizes to generate mitotic chromosomes of appropriate dimensions.

## Results

### A subset of CTCF sites remains bound in mitotic mESC

In the past several years several genomics studies have reported contradictory results on the cell cycle binding dynamics of CTCF, especially during mitosis (Zhang et al. 2019; Owens et al. 2019; Oomen et al. 2019). These studies did not only differ in cell type, but also methodologically, with some cell lines being analyzed by ATAC-seq (HeLa, HFF, U2OS and mESC (Oomen et al. 2019; Owens et al. 2019)), Cut&Run (HeLa (Oomen et al. 2019)) or by ChIP-seq (mESC, C2C12, 3T3, G14E (Owens et al. 2019; Zhang et al. 2019)). Using ChIP-seq, ATAC-seq and Cut&Run, it was shown that in human or mouse differentiated cell lines either all CTCF sites lose binding in mitosis (Zhang et al. 2019; Owens et al. 2019), or show minor signs of mitotic binding (Zhang et al. 2019; Owens et al. 2019); in contrast, ATAC-seq and ChIP-seq revealed extensive mitotic binding of CTCF in mESCs (Owens et al. 2019). It is possible that these differences are the result of the use of different methods. However, these studies do not only differ in genomics techniques and crosslinking conditions, but more notably, they differ in which cell line was used. We hypothesized that reported differences in mitotic retention of CTCF could result from a difference in cell types and species. This would suggest that pluripotent cells can maintain partial CTCF binding in mitosis, whereas somatic cell lines lose CTCF binding in mitosis. To test this directly we compared data obtained with identical experimental methods for pluripotent and somatic cell lines: we compare previous ATAC-seq data generated in pluripotent mouse ESCs (Festuccia et al. 2019) with newly generated ATAC-seq data in differentiated mouse C2C12 cells, using footprinting analyses previously used to show the full eviction of CTCF from human somatic cells in mitosis (Oomen et al. 2019). First, we directly compared previous collections of CTCF binding sites (Owens et al. 2019) that were shown by ChIP-seq to either maintain full binding in mitosis (bookmarked; 10,799 sites), exhibit reduced but detectable binding (reduced; 18,704 sites) or display a complete loss of binding (lost; 22,302 sites) (Owens et al. 2019). By representing ATAC-seq data as V-plots (Zentner and Henikoff 2014; Oomen et al. 2019), we can not only observe accessibility, but also footprints at these specific sets of CTCF sites. When CTCF is bound to chromatin, it will occupy approximately 80 base pairs around its motif. Furthermore, it will push the neighboring nucleosomes away from the motif and into a well-positioned tight array on each side of the motif (Fu et al. 2008; Oomen et al. 2019; Owens et al. 2019).

We can observe these phenomena when we represent ATAC-seq data of non-synchronized mESCs aggregated around CTCF sites that are known to be bound in interphase based on ChIP-seq data (figure 1a). First, the arms of the V cross at approximately 80bp fragment length, the known footprint size of CTCF (Fu et al. 2008). Second, along the arms of the V, dots of enriched signal appear at regular interval (∼280bp, ∼460bp, ∼640bp etc). This ATAC-seq signal indicates the array of well-positioned nucleosomes flanking the bound CTCF motif (Fu et al. 2008). Previously, we found that in differentiated cell lines HeLa, U2OS, and HFF, CTCF sites generally lost accessibility in mitosis (Oomen et al. 2019). When ATAC-seq signal of mitotic differentiated cells was plotted as V-plots, we found that CTCF sites no longer showed enrichment at 80bp fragment length. Instead, the fragment size dropped to much smaller fragment size, suggesting a loss of CTCF binding in mitosis in differentiated cell lines (Oomen et al. 2019).

**Figure 1.**
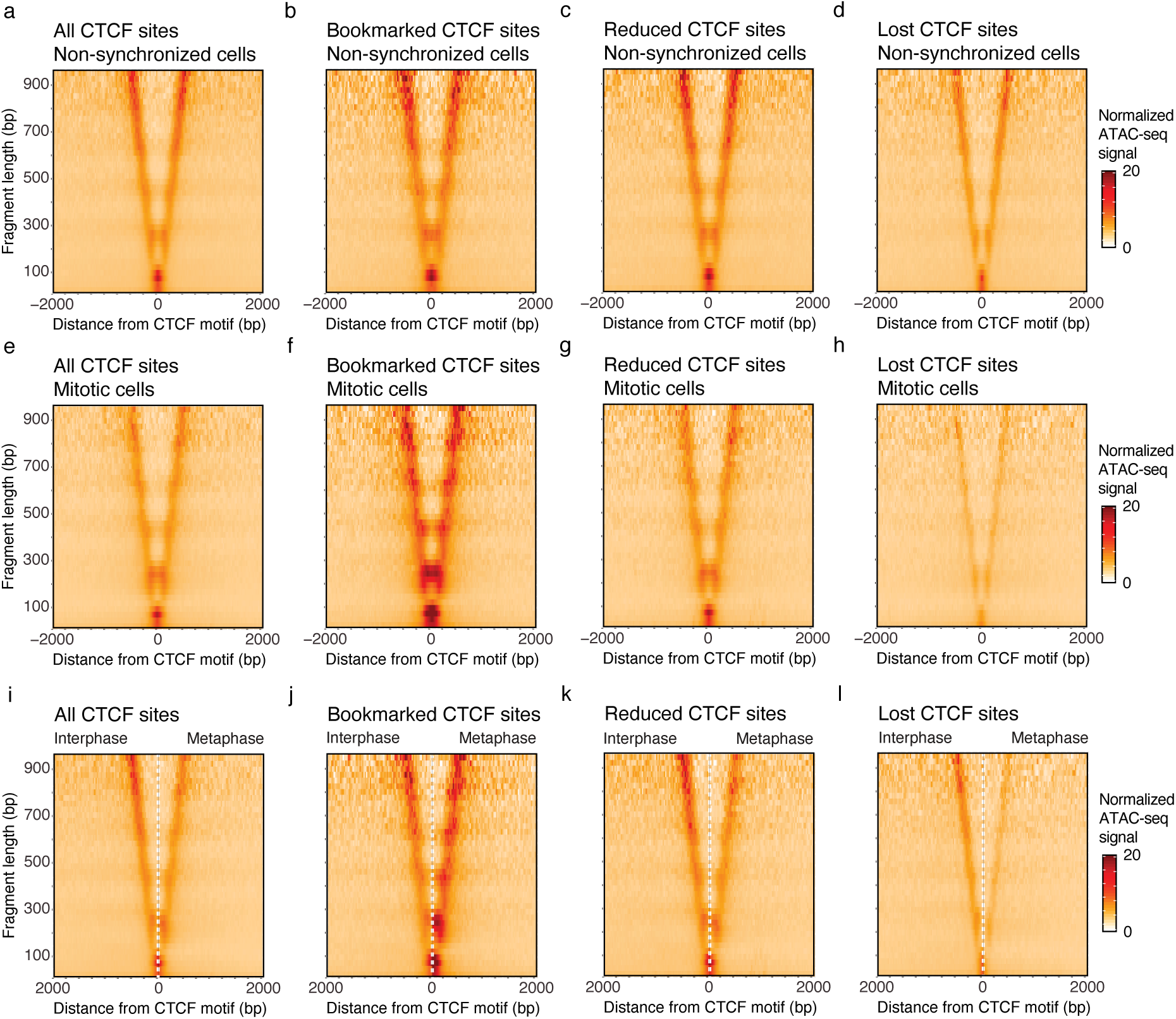
– ATAC-seq data in mESCs show that a group of CTCF motifs remain bound by CTCF in mitosis, whereas other CTCF motifs lose binding. (**a-d**) ATAC-seq data of non-synchronized mESCs represented in V-plots as a pile up on all interphase-bound CTCF sites (51,805 sites total) (a), bookmarked CTCF sites (10,799 sites) (b), CTCF sites with reduced CTCF binding (18,704 sites) (c) and CTCF sites that lose CTCF binding in mitosis (22,302 sites) (d). (**e-h**) ATAC-seq data of mESCs synchronized in mitosis represented in V-plots as a pile up on all interphase-bound CTCF sites (e), bookmarked CTCF sites (f), CTCF sites with reduced CTCF binding (g) and CTCF sites that lose binding in mitosis (h). (**i-l**) Side-by-side comparison of V-plots for non-synchronized and mitotically synchronized cells on all interphase-bound CTCF sites (i), bookmarked CTCF sites (j), reduced CTCF sites (k) and CTCF sites that lose binding in mitosis (l).

When we created V-plots for all interphase-bound CTCF sites in both non-synchronized (figure 1a) and mitotic (figure 1e) mESCs, we observed a less clear picture. First, more accessibility is maintained at CTCF sites in mitotic mESCs compared to differentiated cell lines reported previously (Oomen et al. 2019). When we performed a side-by-side comparison of V-plots of non-synchronized and mitotic cells at the CTCF motif (figure 1i), we observed that the size of the CTCF footprint and the positioning of the nucleosomes along the arms of the V drop down to shorter fragment sizes in mitosis. However, this change is less drastic than what we have observed before in differentiated cell lines. This suggests that there are CTCF sites that maintain mitotic binding as well as CTCF sites that lose binding during mitosis, as we had observed using ChIP-seq (Owens et al. 2019). Indeed, we find that mitotically bookmarked sites (figure 1b, f, j) maintain both ATAC-seq signal and a prominent CTCF footprint in mitosis, indicating high occupancy binding. In contrast, at sites that lost CTCF binding, ATAC-seq signal decreases and the fragment size of the CTCF footprint drops to shorter fragments, confirming the loss of CTCF binding (figure 1d, h, l). ATAC-seq signal at CTCF sites that showed reduced ChIP-seq signal in mitotic mESCs, show a more ambiguous footprint when plotted as V-plots (figure 1c, g, k). This suggest that this category contains sites that are less frequently bound, either in single cells or in the population, an observation that can be extended to lost CTCF sites, which display reduced CTCF footprints in interphase compared to bookmarked sites. Accordingly, the quality of the CTCF motif at lost sites is largely inferior to bookmarked sites (Owens et al. 2019).

To determine whether this partial retention of CTCF along mitotic chromosomes is seen for other mouse cell lines, we performed ATAC-seq in the differentiated mouse cell line C2C12 – a cell line derived from muscle tissue. We find dramatic loss of accessibility of interphase bound CTCF sites in mitosis as well as a loss of binding of CTCF to its motifs when data is represented as V-plots (figure S1a-d). This observation is highly similar to what we previously reported for human differentiated cell lines (Oomen et al. 2019). We note however that the CTCF footprint is not fully lost in mitosis, despite the clear loss of accessibility as observed by loss of signal in the V-plots as well as a reduction in the number of peaks called at CTCF sites (5,827 accessible CTCF motifs in interphase vs 526 in mitosis). This can be observed in the V-plots where we see the remnants of the typical CTCF footprint at 80-100bp fragment size as well as the increase of signal at the CTCF motif itself of very short fragments (>50bp). This could be explained in two ways. (1) It is possible that a small fraction (<10%) of CTCF sites remains bound in mitosis in part of the cell population. Or (2) despite efforts of cell synchronization, a small fraction of the cell population is not fully arrested in prometaphase, but instead have not yet reached full prometaphase arrest or have escaped the mitotic nocodazole arrest.

Taking together these and previous results of ATAC-seq footprinting analyses (Oomen et al. 2019), we confirm that the variable conclusions in the literature regarding the mitotic retention of CTCF are related to cell state differences rather than to species or to technical and analytical differences, with pluripotent cells showing prominent bookmarking of CTCF sites.

### Loss of CTCF-related architectural features in mitosis independently of CTCF binding

The finding that a substantial fraction of CTCF sites maintains binding to mitotic chromosomes in mESCs, raises the question whether CTCF can still function as an architectural protein in mitosis. In interphase cells, chromatin-bound CTCF can block loop extrusion mediated by cohesin (Fudenberg et al. 2016). This results in the formation of TADs and strong interactions between pairs of CTCF sites (CTCF-CTCF loops), which are readily observed by Hi-C (Dixon et al. 2012; Nora et al. 2017; Rao et al. 2017). In mitotic differentiated cell lines, where CTCF binding is lost, no TADs and no CTCF-CTCF or any other site-specific loops are observed (Gibcus et al. 2018; Naumova et al. 2013; Oomen et al. 2019).

Maintained CTCF binding in mitotic mESCs creates the opportunity to study whether mitotic loop extruding machines condensin I and II can be blocked by CTCF, or whether they can shape the characteristic densely packed consecutive loop array unimpeded by bound CTCF. We performed Hi-C on non-synchronized and mitotically synchronized mESCs (figure 2a). In addition to this, we also performed Hi-C on mouse C2C12 cells (figure 2b), which largely loose CTCF binding in mitosis (figure S1) similar to the human differentiated cell lines previously analyzed (Oomen et al. 2019).

**Figure 2.**
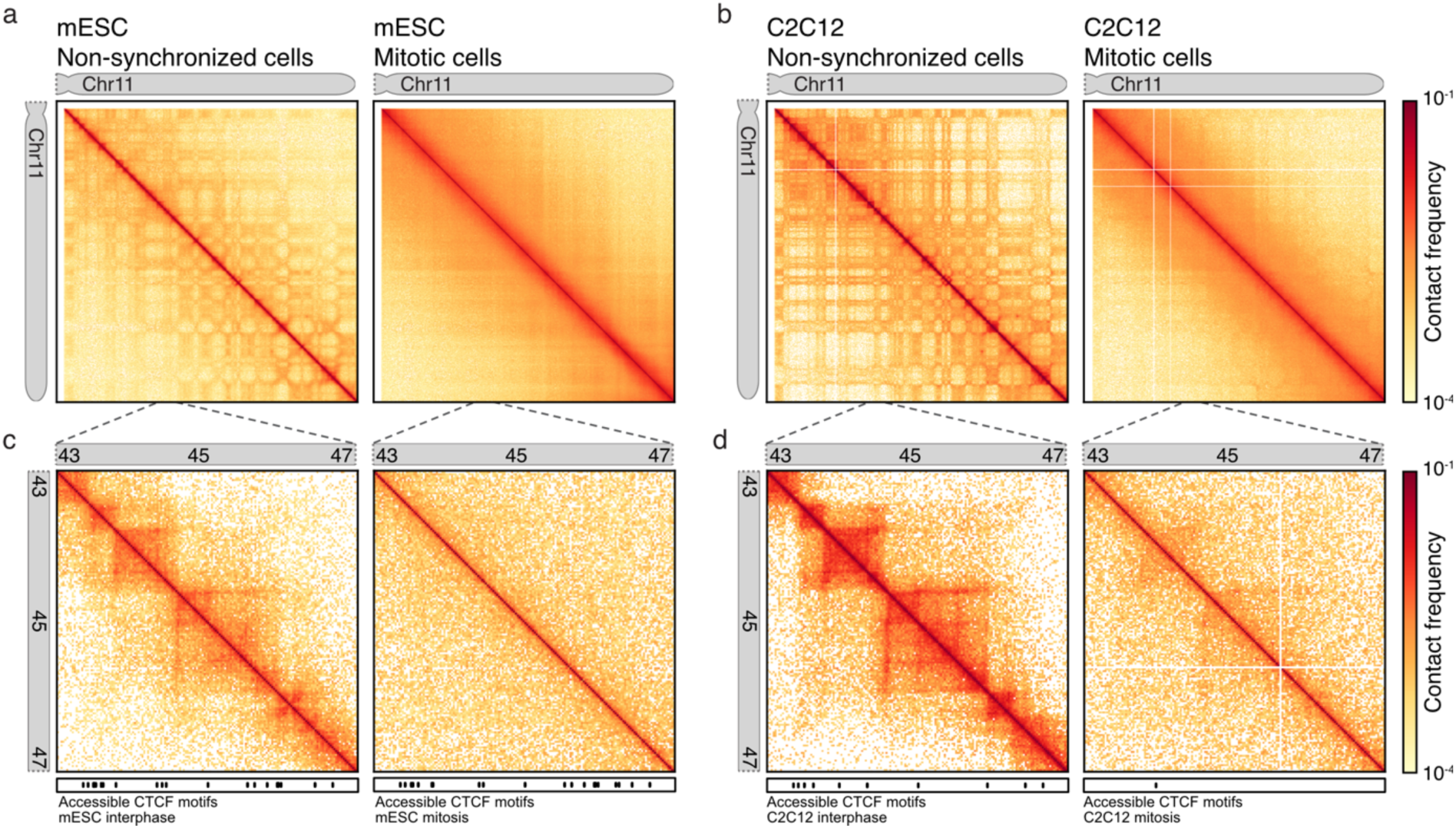
– Hi-C data shows compartments and TADs are lost in both mitotic mESCs and C2C12. (**a-b**) Hi-C heatmap of chr11 at 100kb bins for mESCs (a) and C2C12 (b) non synchronized cells (left panel) and mitotic arrested cells (right panel). (**c-d**) Zoom in Hi-C heatmap of chr11:43,000,000-47,000,000 at 25kb bins for mESC (c) and C2C12 (d) for non-synchronized cells (left panel) and mitotic arrested cells (right panel).

When we plot Hi-C data on a chromosome wide level (figure 2a-b), we observe in interphase cells from both mESC and C2C12 clearly the typical compartment structures, represented as a checkerboard pattern in the heatmaps. Interestingly, the compartment signal in mESCs is much less pronounced compared to C2C12 cells. The strengthening of compartment signal during differentiation has recently been described in human cell lines (Oksuz et al. 2020). When we next examine chromosome-wide heatmaps of mitotic cells, we find that compartments are lost in both C2C12 and mESCs. This is in line with the previous observations in differentiated human cell lines, where compartment signal is lost entirely in mitosis as well (Naumova et al. 2013). We then examined a smaller 4Mb region within chr11 to observe presence or absence of TADs. Whereas in non-synchronized cells, TADs (and domain boundaries positioned at CTCF sites) can be readily observed in both mESCs (figure 2c) and C2C12 cells (figure 2d), in mitosis these structures are lost. A small fraction of contaminating interphase cells can explain the faint compartmental checkerboard and TAD signal that are still detectable (figure 2b).

### Mitotic loop extrusion is not blocked by retained CTCF sites

Next, we set out to analyze CTCF-anchored loops in mitotic mESCs in order to investigate whether mitotic loop extruders condensin I and II are blocked by bound CTCF, which would lead to positioned loops between pairs of CTCF sites and/or domain boundaries at single CTCF site. As described above no compartments and TAD boundaries are detected in mitotic mESCs at individual genomic locations (figure 2). Assessment of the presence of CTCF-dependent loops at specific locations typically requires much deeper sequencing (Akgol Oksuz et al. 2021). To observe these features using our Hi-C datasets, boundaries can be visualized by plotting the aggregate Hi-C signal at and around either single CTCF sites (figure 3a-h), while loops can be visualized by plotting the aggregate Hi-C signal at and around pairwise interactions of CTCF sites (figure 3i-p). In line with the above described ATAC-seq analysis, we used CTCF sites that are categorized based on published ChIP-seq data (Owens et al. 2019) as mitotic bookmarked sites, mitotically reduced sites, and sites that lose CTCF binding in mitosis.

**Figure 3.**
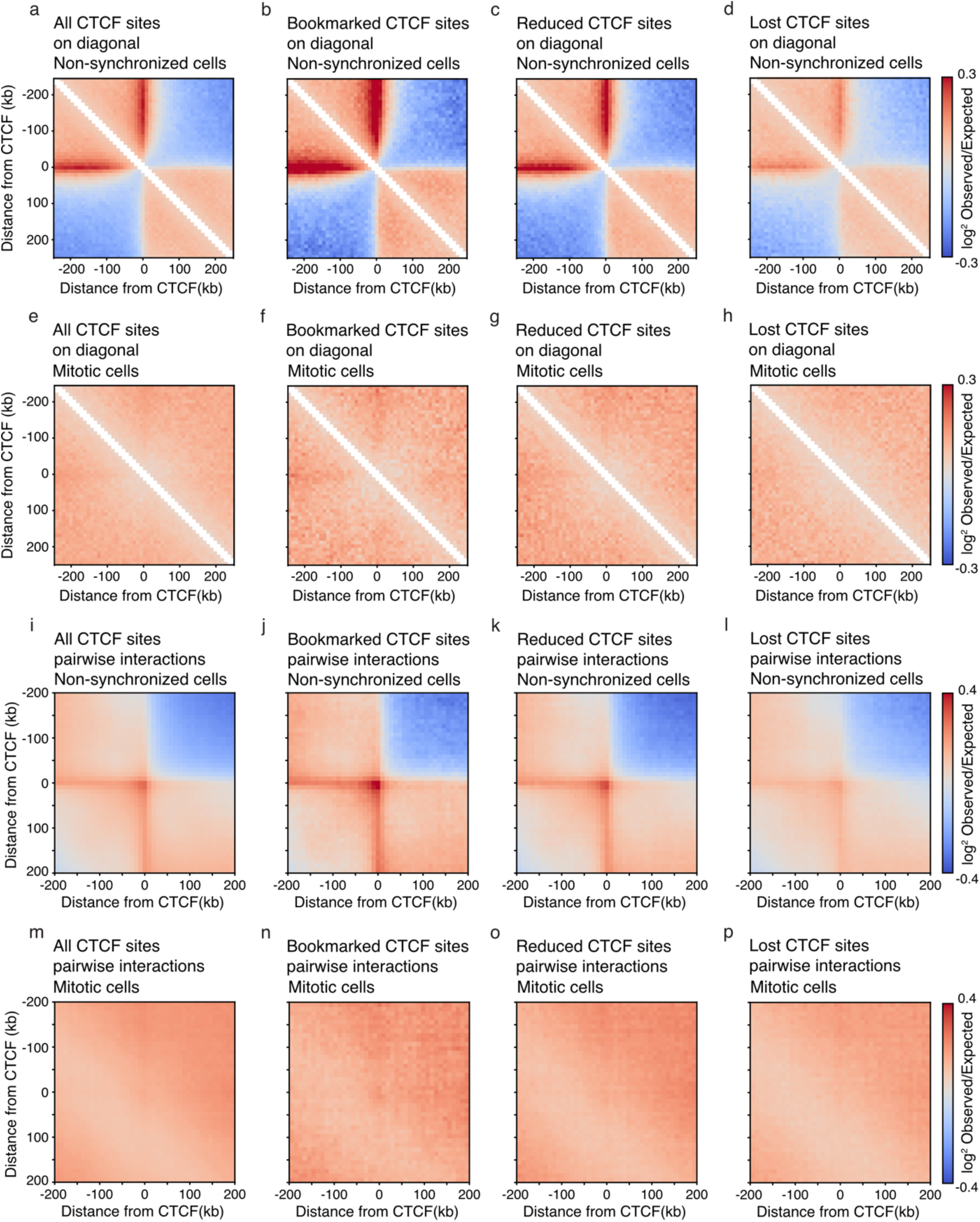
– Hi-C pile-up plots on single and pairwise CTCF sites show that loop extrusion by condensins in mitosis cannot be blocked by bound CTCF. (**a-d**) Aggregate of Hi-C signal binned at 10kb in non-synchronized mESCs on all interphase-bound CTCF sites (a), mitotic bookmarked sites (b), reduced CTCF sites (c), and CTCF sites that lose binding in mitosis (d). (**e-h**) Aggregate of Hi-C signal in mitotic mESCs on all interphase-bound CTCF sites (e), mitotic bookmarked sites (f), reduced CTCF sites (g), and CTCF sites that lose binding in mitosis (h). (**i-p**) Pile up of Hi-C signal in 10kb bins in non-synchronized (i-l) and mitotic (m-p) mESCs of pairwise interactions within 250kb at all interphase bound CTCF sites (i,m), bookmarked CTCF sites (j,n), reduced CTCF sites (k,o) and CTCF sites that lose binding in mitosis (l,p). All CTCF sites are plotted with respect to strand orientation of the motif.

When we aggregate Hi-C signal at and around individual interphase-bound CTCF-sites (i.e., on the diagonal of the Hi-C interaction map), a strong insulating domain boundary can be observed at the center of the pile up plot in interphase cells (figure 3a). This represents the accumulation of insulating potential of CTCF at TAD boundaries, as it reduces the interaction frequency between loci across the bound CTCF site (Dixon et al. 2012; Nora et al. 2017). Insulation can be the result of blocked loop extrusion at CTCF sites and is lost when cohesins are depleted (Rao et al. 2017). Given that blocking of extrusion depends on the orientation of the CTCF motif, a stripe of enriched interactions is detected starting at the CTCF motif and continuing in only one direction. Such directional stripes are hallmarks of blocked loop extrusion and have been reported before (Fudenberg et al. 2016; Vian et al. 2018). Strong evidence for blocked loop extrusion is observed when aggregating Hi-C interactions from non-synchronized cells (mostly interphase cells) on mitotically bookmarked CTCF sites (figure 3b), reduced CTCF sites (figure 3c) and lost CTCF sites (figure 3d). We note that the insulation potential is strongest for bookmarked CTCF sites, compared to that observed at reduced and lost CTCF sites, in line with the differential intensity of CTCF binding at these sites and the presence of motifs of different quality (Owens et al. 2019). Similar to the ATAC-seq experiments described above, Hi-C is performed on a population of cells. Therefore, a possible explanation for the quantitative difference in insulation at these three categories of CTCF sites could be that, in interphase, bookmarked CTCF sites are more likely to be bound by CTCF across the population, whereas reduced and lost CTCF sites are also captured in the unbound state in the population. In contrast, when we plot these same pile-up plots for Hi-C data obtained from mitotic mESCs, we see that all CTCF insulation is lost for each category of CTCF sites (figure 3e-h). This strongly implies that loop extrusion in mitosis is not blocked at sites where CTCF binding is maintained (bookmarked and reduced sites).

Likewise, we can plot the aggregation of Hi-C signal on pairwise CTCF interactions. We curated a list of all possible pairwise interactions between two CTCF sites separated by up to 250 kb. Typically, pairwise CTCF interactions are enriched in Hi-C interaction signal in interphase, as can be observed as a dot in the center of the pile-up plot representing loops between pairs of CTCF sites. Indeed, we see a clear enrichment at pairwise CTCF interactions in non-synchronized mESCs across all categories of CTCF sites (figure 3i-l). This enrichment at pairwise CTCF sites is lost in mitosis for all three categories of CTCF sites (figure 3m-p). Combined these results suggest that although CTCF binding is maintained in mitosis at a substantial fraction of sites in mESCs, CTCF does not have the ability to block mitotic loop extruders condensin I and II and therefore no CTCF-CTCF loops are formed. These results also strongly suggest that by prometaphase there are no extruding cohesin complexes active on the chromosomes, as previously suggested by Smc1 ChIP-seq in nocodazole-arrested mESCs (Owens et al. 2019).

### Mitotic loop sizes differ between species

Hi-C data can be represented as a distance decay plot, where the interaction frequency *P* is plotted as a function of the genomic distance *s*. These *P*(*s*) plots have distinct shapes for both interphase and mitotic chromosomes (Naumova et al. 2013). By calculating the slope of *P*(*s*) and plotting the derivative of contact frequency as a function of genomic distance, the average loop sizes present in interphase and mitosis can be revealed (Abramo et al. 2019; Haarhuis et al. 2017; Gassler et al. 2017; Gibcus et al. 2018; Schwarzer et al. 2017; Polovnikov et al. 2023). Such derivative plots display a characteristic peak around 100-200 kb for interphase cells, and at larger genomic distances for mitotic cells, corresponding to the genomic distance where *P* decays most slowly. This genomic distance is correlated to the average loop size, generated by either cohesins (in interphase), or condensins (in mitosis) (Gassler et al. 2017; Gibcus et al. 2018; Polovnikov et al. 2023).

In addition to any differences between stem cells and differentiated cells, we were interested to study the loop characteristics of different species in interphase and mitosis. We supplemented the Hi-C data generated in this study with data from several studies which included Hi-C data on both non-synchronized and mitotic cells in different species (Naumova et al. 2013; Gibcus et al. 2018; Fitz-James et al. 2020). This enabled the comparison of chicken cells (cell line DT40), human cells (cell line HeLa) and mouse cells (cell lines mESCs, C2C12 and C127). We chose to plot the derivatives of *P*(*s*) for an acrocentric chromosome of similar length to allow for proper comparison between species (chr14 for both mouse and human and chr1 for chicken), although we did not find differences when calculating derivative plots for different chromosomes (figure S2a). In non-synchronized cell populations, Hi-C data from all species, and cell types behaved similarly (figure 4a) with an average interphase loop size of ∼100kb (as highlighted with the arrow in figure 4a). Interestingly, this is not the case for mitotic loops of these different species (figure 4b and zoom in figure 4c). Although there is no difference between the derivative plots of the three mouse cell lines analyzed (mESCs and the differentiated cell lines C2C12 and DT40), a clear difference is observed between mitotic cells of human, mouse, and chicken (Gibcus et al. 2018). All mouse cell lines show an average mitotic loop size of 1-2 megabase (figure 4c, highlighted with circle), whereas human cell line HeLa shows a loop array size of 500-750 kb in mitosis (figure 4c, highlighted with triangle), and chicken cell line DT40 has an average loop size of 200-350 kb (figure 4c, highlighted with star). This suggests a different level of mitotic compaction between the three species.

**Figure 4.**
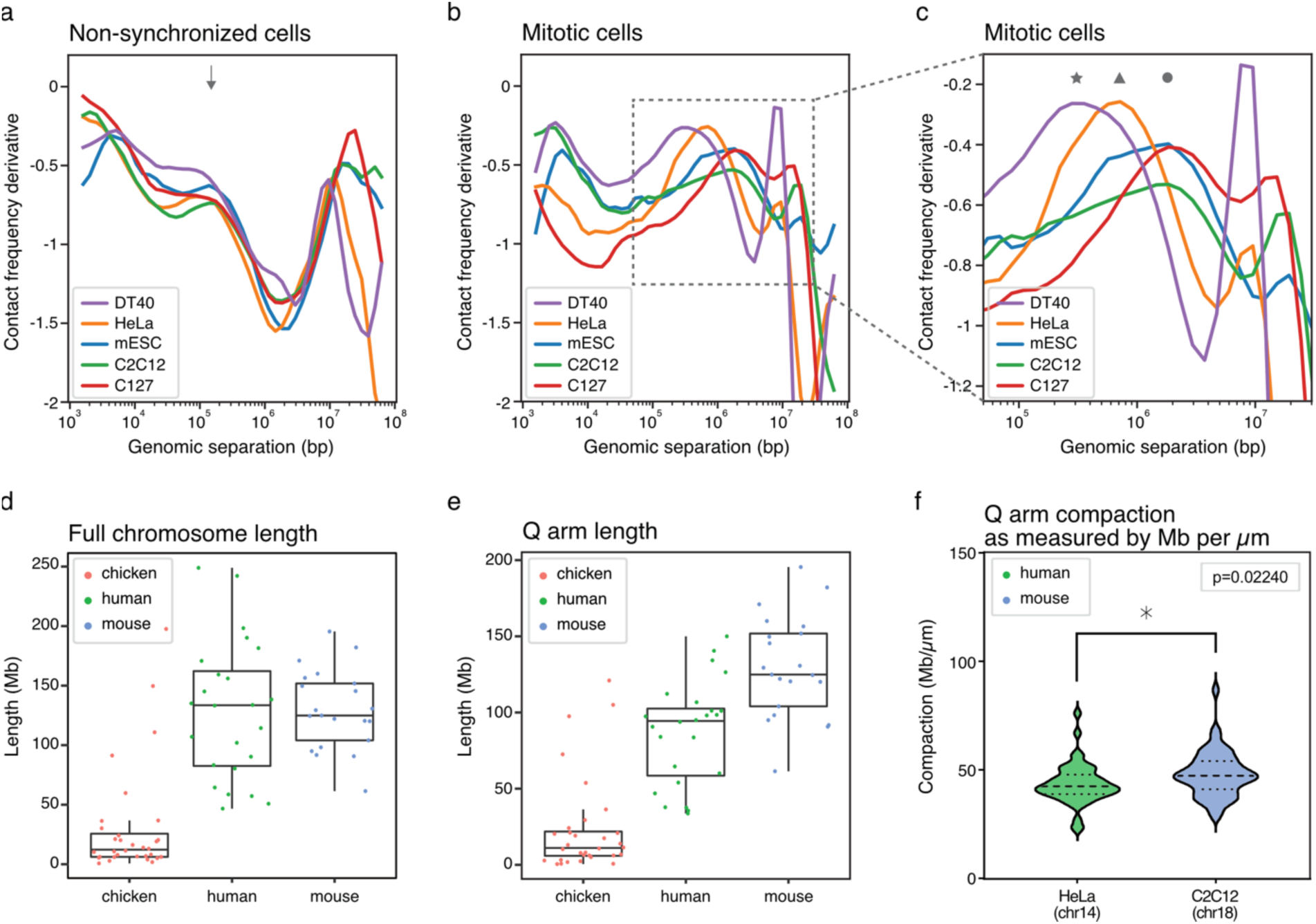
– Mitotic loop arrays species differ in average loop size between species. (**a**) Derivative of *P*(*s*) as a function of genomic separation in non-synchronized chicken cells (DT40, chr1), human cells (HeLa, chr14) and mouse cells (mESCs, C2C12 and C127, chr14). The arrow highlights the average loop size mediated by cohesin in interphase in all cell types and species (**b**) Derivative plots of Hi-C data from chicken cells (DT40, chr1), human cells (HeLa, chr14) and mouse cells (mESCs, C2C12 and C127, chr14) synchronized in mitosis. (**c**) A zoom-in of the derivative plot shown in figure 4b. The star highlights the average loop size observed in mitotic chicken cells, the triangle highlights the average loop size in mitotic human cells and the circle highlights the average loop size in mitotic mouse cells. (**d**) Boxplot of full chromosome lengths in chicken genome (galGal6), human genome (hg38) and mouse genome (mm10). Dots represent individual chromosomes. (**e**) Boxplot of all q-arm lengths in chicken genome (galGal6), human genome (hg38) and mouse genome (mm10). Dots represent individual chromosomes. (**f**) Q-arm compaction as measured by microscopy as Mb/μM in mitotically synchronized HeLa cells (chr14) and C2C12 (chr18). Asterix shows significant difference between arm compaction in mouse and human (n=50, unpaired t-test).

We hypothesized that this difference in average loop sizes could be related to the genomic lengths of chromosomes in the different species. When loops are longer, mitotic chromosomes will become shorter. Possibly, longer chromosomes require a higher level of compaction (shortening along their length), which can be achieved by formation of larger mitotic loops, to ensure proper separation of sister chromatids during anaphase. When we plot all genomic lengths of all chromosomes of the three species (figure 4d), it becomes clear that chicken chromosomes are on average much shorter than human and mouse chromosomes, with a few chromosomes being almost as long as human chromosomes. Mouse and human chromosomes have similar average chromosome length, but the longest mouse chromosome is considerably longer than the longest human chromosome. The centromere is an important region of mitotic chromosomes where the mitotic spindle will attach, which will pull the sister chromatids apart during anaphase (McKinley and Cheeseman 2015). We realized that it is therefore more relevant to plot the length of the longest arm of each chromosome, per definition the q-arm, rather than plotting the full chromosome lengths. Indeed, when we compare the q-arm length between these three species, we find that chicken has very short q-arms with an average length of 11Mb, followed by human chromosomes with an average q-arm length of 94 Mb, and an average q-arm length of 125 Mb for mouse chromosomes (figure 4e). For a given organism loop size is most likely set to ensure that the longest arms are sufficiently compacted. The longest arm in chicken cells is shorter than the longest arm in human cells, and the longest arm in human cell is shorter than the longest arm in mouse. To confirm the hypothesis that loop sizes along mitotic chromosomes are regulated to ensure appropriate shortening of chromosomes, we experimentally measured the q-arm length in mitotic human and mouse cells for two chromosomes of highly similar length by microscopy (chr18 in mouse and chr14 in human, both acrocentric chromosomes with q-arm lengths around 90 Mb). As expected, based on the fact that mitotic loops are larger in mouse (supplementary figure S2), we find that mouse chromosome 18 compacts to a greater extent than human chromosome 14, reflected in a higher megabase per micrometer ratio (figure 4f, supplementary figure S3).

Combined, these results show that in the cell lines we investigated mitotic loop sizes are not related to cell type or differentiation state, but instead differ among species. Moreover, our results suggest that there is a relationship between average genomic q-arm length and the level of mitotic chromosome compaction through modulation of mitotic loop size, as shown using both genomics and microscopy techniques. This ensures even the longest arms are sufficiently compacted to ensure their segregation.

## Discussion

In this study, we set out to explore mitotic chromosome organization in different cell types and vertebrate species. Although mitotic chromosomes are often perceived as universal structures, there are several characteristics that can differ between differentiation state and between species. First, using a single analytical method in side-by-side comparisons, we confirm partial maintenance of CTCF binding in mitotic mESCs and a large eviction in differentiated cells, whether originating from mouse or human (Oomen et al. 2019; Owens et al. 2019). Interestingly, when mESCs are investigated by Hi-C, we observe that no interphase structures are maintained in mitosis despite maintained CTCF binding, suggesting that CTCF does not block mitotic loop extrusion by condensins, and a loss of loop extruding cohesin complexes. Lastly, we investigate whether mitotic chromosomes are differently organized between species. For this analysis, we generated Hi-C data for mouse cell lines and publicly available data for mitotic human and chicken cell lines (Gibcus et al. 2018; Fitz-James et al. 2020). Although further experiments will be necessary, we find that the sizes of mitotic loops are different between species, but do not change between different cell lines of the same organism. Furthermore, our results suggest that mitotic loop size, and therefore the degree of chromosome compaction, are correlated with the average length of the q-arm of chromosomes; a phenomenon that we confirmed by microscopy.

The result that mESCs maintain bookmarking of CTCF binding at a substantial fraction of sites, raises the key question of why it is largely evicted in most, if not all, differentiated cell types displaying condensed chromosomes, including mouse sperm cells in meiosis II (Jung et al. 2017) and mouse oocytes (Wang et al. 2023). CTCF is a C2H2 zinc finger protein, which are canonical transcription factors subject to mitotic phosphorylation and abolishment of their DNA binding capacity (Rizkallah and Hurt 2009; Dephoure et al. 2008; Dovat et al. 2002). Indeed, previous observations showed CTCF is phosphorylated during mitosis (Sekiya et al. 2016). Thus, the eviction of CTCF in mitosis might be the norm, and its retention in mESCs result from a lack of phosphorylation events. Alternatively, it is also possible that the chromatin remodelers associated with CTCF binding, such as SNF2H/L (Wiechens et al. 2016), may be differentially regulated in mitotic mESCs. A second important question raised by our findings is to what extent is mitotic binding by CTCF functional. Recent work has shown that while mitotic CTCF binding correlates with rapidly reactivated genes after mitosis (Zhang et al. 2019; Pelham-Webb et al. 2021; Chervova et al. 2022), the depletion of CTCF at the M/G1 transition affects a minor fraction of its mitotic targets and, especially, those displaying promoter restricted binding (Zhang et al. 2019; Chervova et al. 2022). Nevertheless, correlative studies have suggested that mitotic CTCF binding events are associated with early TAD restoration after mitosis (Pelham-Webb et al. 2021), and the functional depletion of CTCF during M/G1 transition was found associated with a general lack of TAD formation in G1 and the persistence of inappropriate enhancer-promoter contacts (Zhang et al. 2019). Thus, it is possible that mitotic binding events of CTCF, particularly in mESCs, are required for the fidelity of gene regulation more than for transcription levels per se. Interestingly, we note that mouse stem and progenitor cells have a much faster cell cycle compared to many differentiated cell lines (∼12 hours in mESCs vs 24 hours in HeLa cells), which could necessitate fast re-start of transcription initiation upon mitotic exit.

Unfortunately, we have not been able to test our hypotheses on retained mitotic CTCF binding human embryonic stem cells due to our inability to obtain pure populations of living prometaphase-arrested human stem cells. Although we can only speculate about the potential function of maintained CTCF binding upon G1 entry, we did not observe any function related to mitotic chromosome folding by bound CTCF during mitosis. When representing Hi-C data as individual loci or as pileups of Hi-C signal on CTCF sites, we did not find any evidence of TADs or CTCF loops as a result of maintained CTCF binding in mitosis.

Analyzing the average loop length in mitotic mouse cells we noted a much longer length compared to previous studies with human samples. Indeed, analyzing mouse, human and chicken data we could robustly identify species-specific differences in the average length of mitotic loops. It has been shown that mitotic loop arrays are formed by the combined action of condensin I and II, where condensin II mediates loop formation in large loops with several smaller loops inside formed by condensin I (Gibcus et al. 2018). Additionally, the ratio of condensin I and II modulates the level of condensation and the average loop sizes, as has been observed as cell progress from prophase to mitosis (Gibcus et al. 2018), during development in mitotic *Xenopus* chromosomes (Kieserman and Heald 2011), and when mitotic chromosomes are depleted of either condensin I or II (Shintomi and Hirano 2011). We present additional evidence that when chromosomes have longer arms on average, e.g., in mouse as compared to chicken, sister chromatids compact to a greater extent and due to the formation of larger mitotic loops. This process can possibly be mediated by loading different ratios of condensin I and II on mitotic chromosomes, or different absolute levels of condensin (Zhou et al. 2023; Choppakatla et al. 2021). Although it has been described that vertebrate species appear to have different ratios of condensin I and II (Vagnarelli 2012; Ohta et al. 2010; Ono et al. 2003; Hirota et al. 2004; Green et al. 2012), to our knowledge this has not yet been systematically studied in relation to mitotic loop size and chromosome dimensions, with the exception of recent reports in budding yeast and the Xenopus embryo (Kakui et al. 2022; Zhou et al. 2023).

Overall, we find that mitotic chromosomes have characteristics that can differ between cell type identity, differentiation state, and between species. On a detailed level we observe that CTCF binding is effectively maintained at a subset of its targets in mESC during mitosis whereas it is lost in differentiated cell lines. Additionally, our Hi-C analyses directly imply that condensins are not blocked by mitotically bound CTCF, at least in mESC, as we do not observe any remaining CTCF-CTCF loops in mitosis by Hi-C. Our finding that condensins are oblivious to interphase architectural proteins such as CTCF suggests that cell and species-specific differences in retention and bookmarking of such proteins can be tolerated without compromising mitotic chromosome compaction and segregation. On a larger chromosome-wide scale, we observe that the size of mitotic loops can differ between species, which could be correlated to the average length of the chromosome q-arm. Although all vertebrate mitotic chromosomes are folded as an array of loops mediated by condensin I and II, the ratio and absolute levels at which condensins are loaded onto chromosomes could modulate the dimensions of chromosomes and to generate long and thin or short and wide chromosomes.

## Supplemental figures

**Figure S1.**
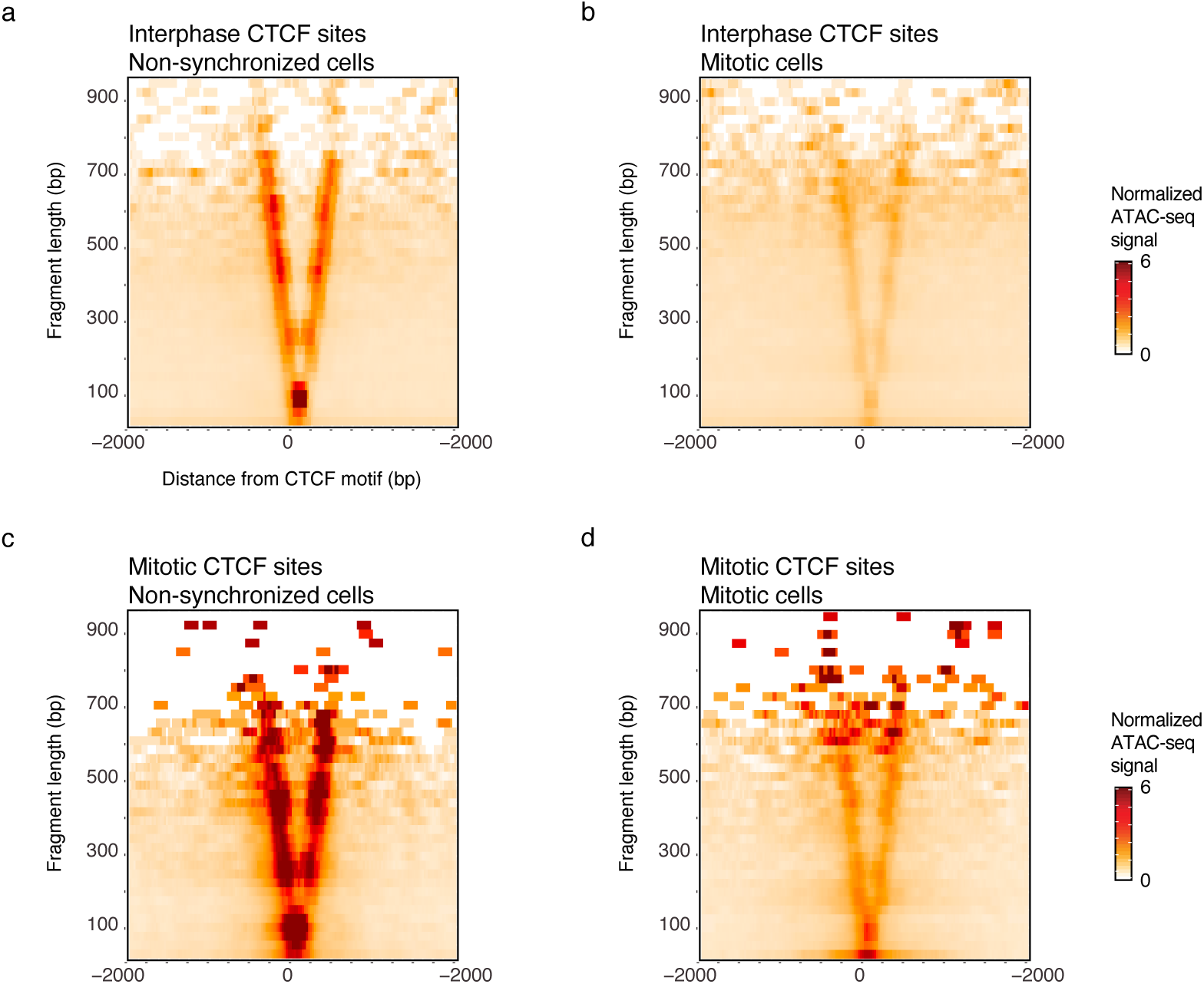
– ATAC-seq data in C2C12 cells show that CTCF binding is largely lost in mitosis. (**a-b**) ATAC-seq data of non-synchronized (a) and mitotically synchronized (b) C2C12 cells represented in V-plots as a pile up on all interphase-bound CTCF sites (5,827 sites total). (**c-d**) ATAC-seq data of non-synchronized (c) and mitotically synchronized (d) C2C12 cells represented in V-plots as a pile up on all mitotic-bound CTCF sites (526 sites total).

**Figure S2.**
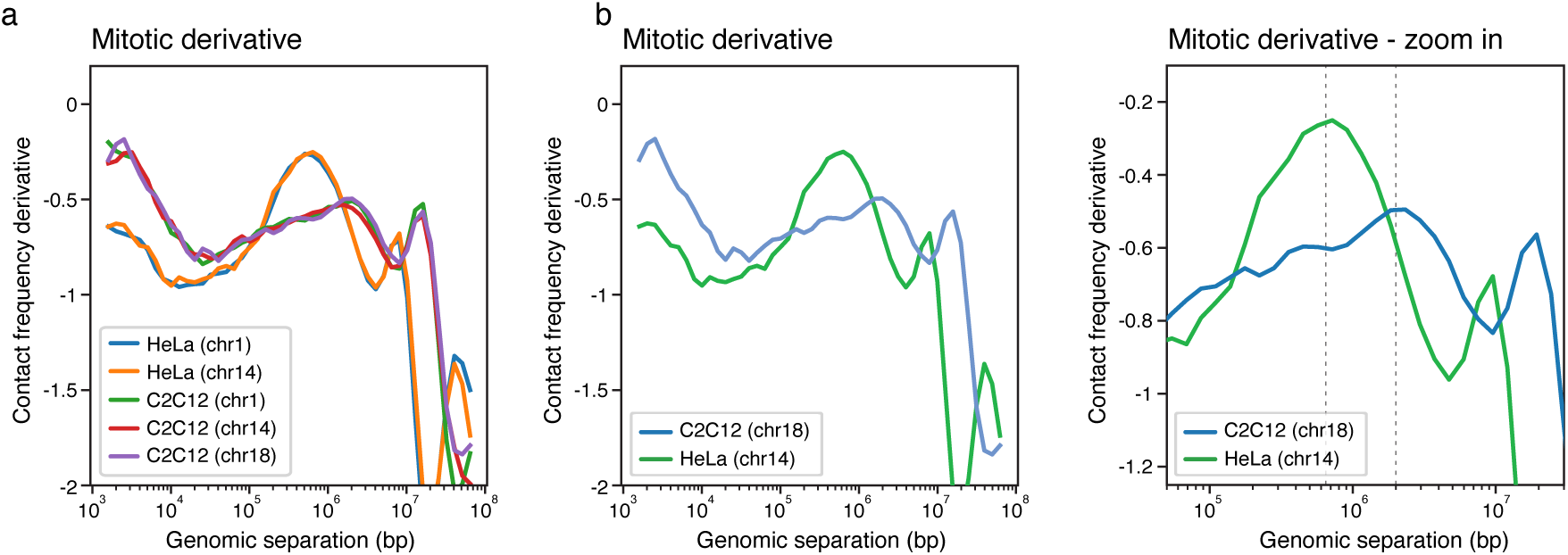
– Mitotic loop sizes for chromosomes investigated by Hi-C. (**a**) Mitotic derivative plots of different chromosomes in HeLa and C2C12 show identical loop sizes across chromosomes (**b**) Derivative plots of Hi-C data from HeLa (chr14) and C2C12 cells (chr18) synchronized in mitosis, which were investigated by microscopy in figure 4f. (**c**) A zoom-in of the derivative plot shown in figure 4b, with dashed lines marking the peak in the derivative at 650kb and 2Mb for human and mouse, and that is correlated with mitotic loop sizes.

**Figure S3.**
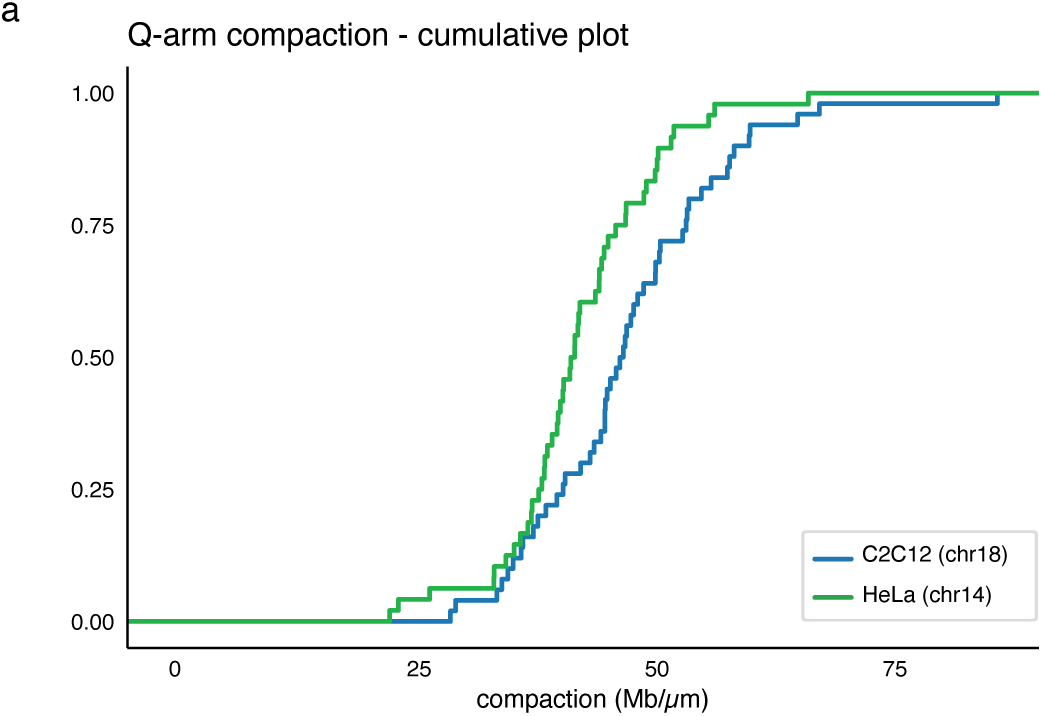
– Cumulative plot of the Q-arm compaction as measured by microscopy. (**a**) Chr14 in mitotically synchronized HeLa cells and chr18 in synchronized C2C12 cells.

## Methods

### Cell culture and synchronization conditions

Mouse embryonic stem cells (E14TG2a) were cultured and synchronized with a 6 hour nocodazole arrest following previous publications (Festuccia et al. 2016, 2019). HeLa and C2C12 cells were cultured in DMEM media supplemented with Glutamax-I, 10% heat-inactivated FBS and penicillin-streptomycin.

C2C12 cells were synchronized with nocodazole arrest (50ng/mL) for 8 hours. Mitotic C2C12 and mES cells were harvested by mitotic shake off. Both mitotic and asynchronous cultures were fixed with 1% formaldehyde and stored at –80°C until processed for Hi-C.

### ATAC-seq

C2C12 cells were cultured as above and arrested in prometaphase using 100ng/mL nocodazole for 12 hours, and mitotic cells harvested by shake-off. The purity of the preparations was assessed by DAPI staining and microscopy and shown to contain 5% of remnant interphase cells. Chromatin accessibility was probed using an adaptation of the ATAC-seq (transposase accessible-chromatin-seq) (Buenrostro et al. 2015). Briefly, 100,000 cells were harvested, washed with PBS. Instead of using lysis buffer to isolate nuclei, cells were pelleted by centrifugation for 5 min at 500g at 4°C, resuspended in 50 μl of transposition reaction mix (25 μl of Tagmentation DNA buffer, 2.5 μl Tagment DNA enzyme (Illumina Tagment DNA TDE1 Enzyme and Buffer Kits, Cat# 20034197) and 22.5 μl nuclease-free H2O) and incubated for 30 min at 37°C with gentle agitation. Reactions were stopped by adding the appropriate volume of Binding Buffer (Qiagen MinElute PCR Kit) and the DNA was purified using the Qiagen MinElute PCR Kit according to manufacturer’s protocol. The purified DNA, eluted in 10 μl, was either stored at – 20°C or used directly for library preparation. ATAC-seq libraries were generated using 10 μl transposed DNA, custom made Illumina barcodes previously described (Buenrostro et al. 2013) and KAPA HiFi HotStart (KapaBiosystems KM2602) for PCR amplification. The number of PCR cycles for PCR amplification was determined using qPCR. Following PCR-amplification, libraries were purified using SPRI beads, using a sample to bead ratio of 1: 1.4. Concentration and fragment size distribution was determined using an Agilent 2200 Tapestation. ATAC-seq libraries were paired-end sequenced on Illumina NextSeq500 using 75 bp paired-end reads in biological duplicates.

### ATAC-seq analysis

ATAC-seq sequencing reads were trimmed to 24bp and aligned to reference genome mm10 using Bowtie2 with a maximum mapping length of 2000bp (Langmead and Salzberg 2012; Buenrostro et al. 2013). Paired-end reads were filtered for mapping quality, mitochondrial reads and PCR duplicates. mESC ATAC-seq data was plotted as V-plots (Zentner and Henikoff 2012) on all interphase bound CTCF motifs (51805 sites) and on CTCF motifs categorized as bookmarked (10799 sites), reduced (18704 sites) or lost (22302 sites) in mitosis as characterized by Owens et al (Owens et al. 2019). V-plots were produced as described (Oomen et al. 2019). To plot V-plots, CTCF motifs were oriented in the same direction. C2C12 ATAC-seq data were analyzed and processed as described in (Oomen et al. 2019). Interphase and mitotic bound CTCF sites were identified when a peak in ATAC-seq data overlapped with a CTCF motif in interphase and/or mitosis.

### Hi-C

Hi-C on mitotic and asynchronous cultures were performed according to previously published protocol (Belaghzal et al. 2017). Briefly cells were fixed and stored as described above. Crosslinked cells were thawed, lysed, and digested with DpnII restriction enzyme overnight at 37°C. Restriction overhangs were filled with biotin-14-dATP supplemented with dTTP, dCTP and dGTP for 4 hours at 23°C, followed by ligation using T4 DNA ligase at 16°C for another 4 hours. Samples were then treated with proteinase K at 65°C overnight. DNA was cleaned up and purified using phenol:chloroform and ethanol precipitation. DNA was sonicated and size selection to average size of 100-350bp using AMpure XB beads, followed by end repair. Samples were enriched for biotin-tagged DNA fragments by pull down using streptavidin beads. After A-tailing, libraries were ligated with indexed Illumina TruSeq sequencing adapters, followed by pcr amplification. Finally, libraries were cleaned up from PCR primers using Ampure XP beads and sequenced using paired-end 50bp sequencing on an Illumina HiSeq 4000.

### Hi-C mapping and downstream analysis

Hi-C sequencing files were mapped to reference genomes hg38 (HeLa data), mm10 (C2C12, mESC and C127 data) and galGal6 (DT40 data) using publicly available distiller-nf mapping pipeline (https://github.com/mirnylab/distiller-nf) and downstream analysis tools pairtools (https://github.com/mirnylab/pairtools) and cooltools (https://github.com/mirnylab/cooltools). Briefly, reads were mapped using bwa-mem, pcr duplicates were removed and reads were filtered for mapping quality. Distance decay and derivative plots created using cooltools code by calculating contact frequency (P) as a function of genomic distance (*s*) using valid pairs. For further downstream analysis, interactions were binned in matrices at a range of different resolutions using cooler (Abdennur and Mirny 2019). Iterative balancing was applied to all matrices, while ignoring the first two bins from the diagonal (Imakaev et al. 2012). Pile up plots at single CTCF sites and pairwise CTCF interactions were produced using observed over expected signal binned at 10kb. Pairwise CTCF sites for pile up plots were predicted by pairing all CTCF sites within 250kb on the same chromosome within the CTCF category (CTCF sites bookmarked in mitosis, reduced in mitosis or lost in mitosis) following curation by N.O and P.N. Directionality of the CTCF motifs were taken into account and all motifs were orientated in the same direction.

### Mitotic chromosome spreads and chromosome labelling for imaging

Asychronous HeLa or C2C12 cultures were incubated in 0.1 ug/mL colcemid (Sigma Aldrich, 10295892001) for 2 hours. Cells were collected after trypsinization, spun down at 4°C at 1000g for 10 minutes and all but 500 uL media removed. Cells were then resuspended in the remaining media, and 5 mL prewarmed (37°C) 75mM KCl added dropwise. Cells were swollen at 37°C for 10 minutes, then fixed in freshly made ice cold 3:1 methanol acetic acid. Aliquots of the fixed samples were then dropped on slides, and the slides set, chromosome side up, over a beaker with 70°C –80°C distilled water for 30 seconds. Slides were then air-dried and incubated at 37°C overnight prior to using for DNA-FISH experiments. To identify HeLa S3 chromosome 14 and C2C12 chromosome 18, custom Atto 565-labeled MyTags libraries (Arbor Biosciences/Daicel) were used to stain mitotic chromosomes spreads (HeLa— chr14:100674834-100852919; C2C12—chr18:88639179-88816381). Centromeres were labeled with the pan-centromeric probe CENP-B-Cy5 (PNA Bio, F3005). After DNA FISH and CENP-B probe labeling, slides were stained in 300 nM 4′,6-diamidino-2-phenylindole (DAPI, ThermoFisher Scientific, D1306) and mounted in ProLong Diamond antifade mountant (Invitrogen, P36965).

### Confocal Fluorescence Imaging

Confocal images were acquired on a Leica SP8 spectral confocal microscope (housed in UMass Chan’s Sanderson Center for Optical Experimentation, SCOPE; RRID: SCR_022721) equipped with a 63×/1.40 NA PL Apo CS2 oil immersion lens (Leica); 405 nM and 638 nM Diode lasers and 552 nM OPSL laser; and sCMOS cameras (pco.edge). For HeLa chromosomes, the spectral detector settings used were PMT 410nm-560nm (405 laser), HyD2 560-633nm (552 laser), and HyD3 643-783 (638 laser). For C2C12 chromosomes, the spectral detector settings used were PMT 410nm-575nm (405 laser), HyD2 557-778nm (552 laser), and HyD3 643-783 (638 laser). Pixel size was 24 nm, frame size was 1024×1024, and zoom was 7.6X. Image stacks with 0.3 μm thick z sections were acquired using immersion oil with a refractive index of 1.518. After image acquisition, Lightening deconvolution was applied to each image stack.

### Image Analysis

Chromatid length was measured using Fiji (Schindelin et al. 2012). Image stacks were projected into maximum intensity Z-projections. Hela chromosome 14 and C2C12 chromosome 18 were identified by FISH DNA probe staining, and one sister chromatid was measured for length (from the end of the arm to the beginning of the centromere stained by CENP-B). 50 C2C12 chromatids and 49 HeLa chromatids were measured. Length measurements were analyzed in GraphPad Prism 9.5.1, using an unpaired t-test.

## Code availability

Hi-C mapping pipeline distiller-nf is available on Github: https://github.com/mirnylab/distiller-nf. Downstream analysis tools pairtools and cooltools are available through https://github.com/mirnylab/pairtools and https://github.com/mirnylab/cooltools. Code used for analysis of ATAC-seq data can be found at Github: https://github.com/dekkerlab/CTCF_in_mitosis_GR_2018.

## Data availability

All sequencing data will be available in GEO upon publication. Microscopy data will be available in the BioStudies database (https://www.ebi.ac.uk/biostudies/) upon publication.

## Publicly available data used in this study

In addition to the Hi-C data that was generated for this study, we use several ATAC-seq and Hi-C datasets that are publicly available on the gene expression omnibus (GEO). ATAC-seq data in mESC (Festuccia et al. 2019) is available under accession number GSE122589. Hi-C data in asynchronous and mitotic DT40 cells and HeLa are available under GSE102740 (Gibcus et al. 2018) and Hi-C data of mouse cell line C127 under GSE149677 (Fitz-James et al. 2020).

## Author contributions

MEO, PN and JD conceived and designed the project. TP and AMP cultured and synchronized mESC cells and MEO cultured and synchronized C2C12 cells for Hi-C experiments. IG performed ATAC-seq in C2C12 cells. MEO performed all Hi-C experiments. MEO analyzed all newly generated and published Hi-C and ATAC-seq datasets in this study with input from JD and PN. ANF performed all microscopy experiments and analysis. MEO and JD wrote the manuscript with input from all authors.

## Acknowledgements

We thank all current and former members of the Dekker lab and the Navarro lab for helpful discussions and suggestions, in particular Johan Gibcus, Bastiaan Dekker and Nick Owens. We thank Eugenio Mattei for advice on computational analyses. We thank Dr. Christina Baer for thoughtful discussions on confocal imaging. This work was supported by a grant from the National Institutes of Health (HG003143 to J.D.) and by the European Research Council (ERC-CoG-2017 BIND to P.N). J.D. is an investigator of the Howard Hughes Medical Institute.

## Notes

### Competing Interest Statement

The authors have declared no competing interest.

## References

1. Abdennur N, Mirny LA. 2019. Cooler: scalable storage for Hi-C data and other genomically labeled arrays. Bioinformatics 36: 311–316.

2. Abramo K, Valton AL, Venev S V., Ozadam H, Fox AN, Dekker J. 2019. A chromosome folding intermediate at the condensin-to-cohesin transition during telophase. Nat Cell Biol 21: 1393–1402. 10.1038/s41556-019-0406-2.

3. Akgol Oksuz B, Yang L, Abraham S, Venev S V., Krietenstein N, Parsi KM, Ozadam H, Oomen ME, Nand A, Mao H, et al. 2021. Systematic evaluation of chromosome conformation capture assays. Nat Methods 18: 1046–1055.

4. Batty P, Gerlich DW. 2019. Mitotic Chromosome Mechanics: How Cells Segregate Their Genome. Trends Cell Biol **xx**: 1–10.

5. Belaghzal H, Dekker J, Gibcus JH. 2017. Hi-C 2.0: An optimized Hi-C procedure for high-resolution genome-wide mapping of chromosome conformation. Methods 123: 56–65.

6. Belmont AS. 2006. Mitotic chromosome structure and condensation. Curr Opin Cell Biol 18: 632–638.

7. Buenrostro JD, Giresi PG, Zaba LC, Chang HY, Greenleaf WJ. 2013. Transposition of native chromatin for fast and sensitive epigenomic profiling of open chromatin, DNA-binding proteins and nucleosome position. Nat Methods 10: 1213–8. http://www.pubmedcentral.nih.gov/articlerender.fcgi?artid=3959825&tool=pmcentrez&rendertype=ab stract (Accessed July 11, 2014).

8. Buenrostro JD, Wu B, Chang HY, Greenleaf WJ. 2015. ATAC-seq: A method for assaying chromatin accessibility genome-wide. Curr Protoc Mol Biol 2015: 21.29.1–21.29.9.

9. Chervova A, Festuccia N, Altamirano-Pacheco L, Dubois A, Navarro P. 2022. A gene subset requires CTCF bookmarking during the fast post-mitotic reactivation of mouse ES cells. EMBO Rep.

10. Choppakatla P, Dekker B, Cutts EE, Vannini A, Dekker J, Funabiki H. 2021. Linker histone H1.8 inhibits chromatin binding of condensins and DNA topoisomerase II to tune chromosome length and individualization. Elife 10. https://elifesciences.org/articles/68918.

11. Dekker J, Mirny L. 2016. The 3D Genome as Moderator of Chromosomal Communication. Cell 164: 1110–1121.

12. Dekker J, Rippe K, Dekker M, Kleckner N. 2002. Capturing chromosome conformation. Science 295: 1306–11.

13. Dephoure N, Zhou C, Villen J, Beausoleil SA, Bakalarski CE, Elledge SJ, Gygi SP. 2008. A quantitative atlas of mitotic phosphorylation. Proceedings of the National Academy of Sciences 105: 10762– 10767. http://www.pnas.org/cgi/doi/10.1073/pnas.0805139105.

14. Dixon JR, Selvaraj S, Yue F, Kim A, Li Y, Shen Y, Hu M, Liu JS, Ren B. 2012. Topological domains in mammalian genomes identified by analysis of chromatin interactions. Nature 485: 376–80.

15. Dostie J, Richmond TA, Arnaout RA, Selzer RR, Lee WL, Honan TA, Rubio ED, Krumm A, Lamb J, Nusbaum C, et al. 2006. Chromosome Conformation Capture Carbon Copy (5C): a massively parallel solution for mapping interactions between genomic elements. Genome Res 16: 1299–309.

16. Dovat S, Ronni T, Russell D, Ferrini R, Cobb BS, Smale ST. 2002. A common mechanism for mitotic inactivation of C2H2 zinc finger DNA-binding domains. Genes Dev 16: 2985–2990.

17. Earnshaw WC, Laemmli UK. 1983. Architecture of metaphase chromosomes and chromosome scaffolds. Journal of Cell Biology 96: 84–93.

18. Erdel F, Rippe K. 2018. Formation of Chromatin Subcompartments by Phase Separation. Biophys J 114: 2262–2270.

19. Fazzio TG, Panning B. 2010. Condensin complexes regulate mitotic progression and interphase chromatin structure in embryonic stem cells. Journal of Cell Biology 188: 491–503.

20. Festuccia N, Dubois A, Vandormael-Pournin S, Gallego Tejeda E, Mouren A, Bessonnard S, Mueller F, Proux C, Cohen-Tannoudji M, Navarro P. 2016. Mitotic binding of Esrrb marks key regulatory regions of the pluripotency network. Nat Cell Biol 18.

21. Festuccia N, Owens N, Papadopoulou T, Gonzalez I, Tachtsidi A, Vandoermel-Pournin S, Gallego E, Gutierrez N, Dubois A, Cohen-Tannoudji M, et al. 2019. Transcription factor activity and nucleosome organization in mitosis. Genome Res 29: 250–260. http://genome.cshlp.org/lookup/doi/10.1101/gr.243048.118.

22. Fitz-James MH, Tong P, Pidoux AL, Ozadam H, Yang L, White SA, Dekker J, Allshire RC. 2020. Large domains of heterochromatin direct the formation of short mitotic chromosome loops. Elife 9: 1–28.

23. Flemming W. 1878. Zur Kenntnis der Zelle und ihrer Teilung-Erscheinungen. Schr Nat Wiss Ver Schlesw-Holst 3: 23–27.

24. Fu Y, Sinha M, Peterson CL, Weng Z. 2008. The insulator binding protein CTCF positions 20 nucleosomes around its binding sites across the human genome. PLoS Genet 4.

25. Fudenberg G, Imakaev M, Lu C, Goloborodko A, Abdennur N, Mirny LA. 2016. Formation of Chromosomal Domains by Loop Extrusion. Cell Rep 1–12. http://biorxiv.org/content/early/2015/08/14/024620.abstract.

26. Gassler J, Brandão HB, Imakaev M, Flyamer IM, Ladstätter S, Bickmore WA, Peters J-M, Mirny LA, Tachibana K. 2017. A mechanism of cohesin-dependent loop extrusion organizes zygotic genome architecture. EMBO J 36: 3600–3618. http://www.ncbi.nlm.nih.gov/pubmed/29217590%0Ahttp://www.pubmedcentral.nih.gov/articlerender.fcgi?artid=PMC5730859-.

27. Gibcus JH, Samejima K, Goloborodko A, Samejima I, Naumova N, Nuebler J, Kanemaki MT, Xie L, Paulson JR, Earnshaw WC, et al. 2018. A pathway for mitotic chromosome formation. Science (1979) 359: eaao6135. http://www.sciencemag.org/lookup/doi/10.1126/science.aao6135.

28. Green LC, Kalitsis P, Chang TM, Cipetic M, Kim JH, Marshall O, Turnbull L, Whitchurch CB, Vagnarelli P, Samejima K, et al. 2012. Contrasting roles of condensin I and condensin II in mitotic chromosome formation. J Cell Sci 125: 1591–1604.

29. Haarhuis JHI, Van Der Weide RH, Blomen VA, Brummelkamp TR, De Wit E, Rowland Correspondence BD, Nl EDWDW, Nl RR. 2017. The Cohesin Release Factor WAPL Restricts Chromatin Loop Extension. 693–707.

30. Hirota T, Gerlich D, Koch B, Ellenberg J, Peters JM. 2004. Distinct functions of condensin I and II in mitotic chromosome assembly. J Cell Sci 117: 6435–6445.

31. Hsiung CC, Morrissey CS, Udugama M, Frank CL, Keller C a, Baek S, Giardine B, Crawford GE, Sung M, Hardison RC, et al. 2015. Genome accessibility is widely preserved and locally modulated during mitosis. 1–29.

32. Imakaev M, Fudenberg G, McCord RP, Naumova N, Goloborodko A, Lajoie BR, Dekker J, Mirny LA. 2012. Iterative correction of Hi-C data reveals hallmarks of chromosome organization. Nat Methods 9: 999–1003.

33. Jung YH, Sauria MEG, Lyu X, Cheema MS, Ausio J, Taylor J, Corces VG. 2017. Chromatin States in Mouse Sperm Correlate with Embryonic and Adult Regulatory Landscapes. Cell Rep 18: 1366– 1382. http://linkinghub.elsevier.com/retrieve/pii/S2211124717300712.

34. Kakui Y, Barrington C, Kusano Y, Thadani R, Fallesen T, Hirota T, Uhlmann F. 2022. Chromosome arm length, and a species-specific determinant, define chromosome arm width. Cell Rep 41: 111753.

35. Kieserman EK, Heald R. 2011. Mitotic chromosome size scaling in Xenopus. Cell Cycle 10: 3863–3870.

36. Langmead B, Salzberg SL. 2012. Fast gapped-read alignment with Bowtie 2. Nat Methods 9: 357–9.

37. Lieberman-Aiden E, van Berkum NL, Williams L, Imakaev M, Ragoczy T, Telling A, Amit I, Lajoie BR, Sabo PJ, Dorschner MO, et al. 2009. Comprehensive mapping of long-range interactions reveals folding principles of the human genome. Science 326: 289–93.

38. Lupianez DG, Kraft K, Heinrich V, Krawitz P, Brancati F, Klopocki E, Horn D, Kayserili H, Opitz JM, Laxova R, et al. 2015. Disruptions of topological chromatin domains cause pathogenic rewiring of gene-enhancer interactions. Cell 161: 1012–1025.

39. Marsden MPF, Laemmli UK. 1979. Metaphase chromosome structure: Evidence for a radial loop model. Cell 17: 849–858.

40. McKinley KL, Cheeseman IM. 2015. The molecular basis for centromere identity and function. Nat Rev Mol Cell Biol.

41. Michieletto D, Orlandini E, Marenduzzo D. 2016. Polymer model with epigenetic recoloring reveals a pathway for the de novo establishment and 3D organization of chromatin domains. Phys Rev X 6: 1–15.

42. Naumova N, Imakaev M, Fudenberg G, Zhan Y, Lajoie BR, Mirny L a, Dekker J. 2013. Organization of the mitotic chromosome. Science 342: 948–53. http://www.pubmedcentral.nih.gov/articlerender.fcgi?artid=4040465&tool=pmcentrez&rendertype=abstract (Accessed July 15, 2014).

43. Nora EP, Goloborodko A, Valton A-L, Gibcus JH, Uebersohn A, Abdennur N, Dekker J, Mirny LA, Bruneau BG. 2017. Targeted Degradation of CTCF Decouples Local Insulation of Chromosome Domains from Genomic Compartmentalization. Cell 169: 930–944.e22.

44. Nora EP, Goloborodko A, Valton A-L, Gibcus JH, Uebersohn A, Abdennur N, Dekker J, Mirny LA, Bruneau BG. 2016. Targeted degradation of CTCF decouples local insulation of chromosome domains from higher-order genomic compartmentalization. bioRxiv.

45. Nora EP, Lajoie BR, Schulz EG, Giorgetti L, Okamoto I, Servant N, Piolot T, van Berkum NL, Meisig J, Sedat J, et al. 2012. Spatial partitioning of the regulatory landscape of the X-inactivation centre. Nature 485: 381–5.

46. Nuebler J, Fudenberg G, Imakaev M, Abdennur N, Mirny LA. 2018. Chromatin organization by an interplay of loop extrusion and compartmental segregation. Proc Natl Acad Sci U S A 196261.

47. Ohta S, Bukowski-Wills J-C, Sanchez-Pulido L, Alves F de L, Wood L, Chen ZA, Platani M, Fischer L, Hudson DF, Ponting CP, et al. 2010. The Protein Composition of Mitotic Chromosomes Determined Using Multiclassifier Combinatorial Proteomics. Cell 142: 810–821.

48. Oksuz BA, Yang L, Abraham S, Venev S V, Krietenstein N. 2020. Systematic evaluation of chromosome conformation capture assays. 0–42.

49. Ono T, Losada A, Hirano M, Myers MP, Neuwald AF, Hirano T. 2003. Differential contributions of condensin I and condensin II to mitotic chromosome architecture in vertebrate cells. Cell 115: 109– 121.

50. Oomen ME, Hansen AS, Liu Y, Darzacq X, Dekker J. 2019. CTCF sites display cell cycle–dependent dynamics in factor binding and nucleosome positioning. Genome Res 1–14.

51. Owens N, Papadopoulou T, Festuccia N, Tachtsidi A, Gonzalez I, Dubois A, Vandormael-Pournin S, Nora EP, Bruneau BG, Cohen-Tannoudji M, et al. 2019. CTCF confers local nucleosome resiliency after dna replication and during mitosis. Elife 8: 1–26.

52. Pelham-Webb B, Polyzos A, Wojenski L, Kloetgen A, Li J, Di Giammartino DC, Sakellaropoulos T, Tsirigos A, Core L, Apostolou E. 2021. H3K27ac bookmarking promotes rapid post-mitotic activation of the pluripotent stem cell program without impacting 3D chromatin reorganization. Mol Cell 1–17. http://www.ncbi.nlm.nih.gov/pubmed/33730542.

53. Polovnikov KE, Brandão HB, Belan S, Slavov B, Imakaev M, Mirny LA. 2023. Crumpled Polymer with Loops Recapitulates Key Features of Chromosome Organization. Phys Rev X 13: 041029. https://link.aps.org/doi/10.1103/PhysRevX.13.041029.

54. Rao SSP, Huang S-C, Hilaire BGS, Engreitz JM, Perez EM, Kieffer-Kwon K-R, Sanborn AL, Johnstone SE, Bascom GD, Bochkov ID, et al. 2017. Cohesin Loss Eliminates All Loop Domains. Cell 171: 305–320.e24. http://www.cell.com/cell/fulltext/S0092-8674(17)31120-0.

55. Rao SSP, Huntley MH, Durand NC, Stamenova EK, Bochkov ID, Robinson JT, Sanborn AL, Machol I, Omer AD, Lander ES, et al. 2014. A 3D Map of the Human Genome at Kilobase Resolution Reveals Principles of Chromatin Looping. Cell 159: 1665–1680.

56. Rizkallah R, Hurt MM. 2009. Regulation of the Transcription Factor YY1 in Mitosis through Phosphorylation of Its DNA-binding Domain ed. A.G. Matera. Mol Biol Cell 20: 4766–4776. http://www.molbiolcell.org/doi/10.1091/mbc.e09-04-0264.

57. Sanborn AL, Rao SSP, Huang S-C, Durand NC, Huntley MH, Jewett AI, Bochkov ID, Chinnappan D, Cutkosky A, Li J, et al. 2015. Chromatin extrusion explains key features of loop and domain formation in wild-type and engineered genomes. Proceedings of the National Academy of Sciences.

58. Schindelin J, Arganda-Carreras I, Frise E, Kaynig V, Longair M, Pietzsch T, Preibisch S, Rueden C, Saalfeld S, Schmid B, et al. 2012. Fiji: An open-source platform for biological-image analysis. Nat Methods 9: 676–682.

59. Schwarzer W, Abdennur N, Goloborodko A, Pekowska A, Fudenberg G, Loe-Mie Y, Fonseca NA, Huber W, Haering CH, Mirny L, et al. 2017. Two independent modes of chromatin organization revealed by cohesin removal. Nature 551: 51–56.

60. Sekiya T, Murano K, Kato K, Kawaguchi A, Nagata K. 2016. Mitotic phosphorylation of CCCTC-binding factor (CTCF) reduces its DNA binding activity. FEBS Open Bio.

61. Shintomi K, Hirano T. 2011. The relative ratio of condensing I to II determines chromosome shapes. Genes Dev 25: 1464–1469.

62. Smith EM, Lajoie BR, Jain G, Dekker J. 2016. Invariant TAD Boundaries Constrain Cell-Type-Specific Looping Interactions between Promoters and Distal Elements around the CFTR Locus. Am J Hum Genet 98: 185–201.

63. Vagnarelli P. 2012. Mitotic chromosome condensation in vertebrates. Exp Cell Res 318: 1435–41.

64. Valton AL, Dekker J. 2016. TAD disruption as oncogenic driver. Curr Opin Genet Dev 36: 34–40.

65. Vian L, Pękowska A, Rao SSP, Kieffer-Kwon KR, Jung S, Baranello L, Huang SC, El Khattabi L, Dose M, Pruett N, et al. 2018. The Energetics and Physiological Impact of Cohesin Extrusion. Cell 173: 1165–1178.e20.

66. Wang F, Higgins JMG. 2013. Histone modifications and mitosis: countermarks, landmarks, and bookmarks. Trends Cell Biol 23: 175–84.

67. Wang W, Gao R, Yang D, Ma M, Zang R, Wang X, Chen C, Chen J, Kou X, Zhao Y, et al. 2023. ADNP Modulates SINE B2-Derived CTCF-Binding Sites during Blastocyst Formation in Mouse 2. BioRxiv. 10.1101/2023.11.24.567719.

68. Wiechens N, Singh V, Gkikopoulos T, Schofield P, Rocha S, Owen-Hughes T. 2016. The Chromatin Remodelling Enzymes SNF2H and SNF2L Position Nucleosomes adjacent to CTCF and Other Transcription Factors. PLoS Genet 12: e1005940. http://www.ncbi.nlm.nih.gov/pubmed/27019336.

69. Wit E De, Vos ESM, Holwerda SJB, Valdes-quezada C, Verstegen MJAM, Teunissen H, Splinter E, Wijchers PJ, Krijger PHL, Laat W De. 2015. CTCF Binding Polarity Determines Chromatin Looping. Mol Cell 60: 1–9.

70. Zentner GE, Henikoff S. 2014. High-resolution digital profiling of the epigenome. Nat Rev Genet 15: 814– 827. http://www.nature.com/doifinder/10.1038/nrg3798.

71. Zentner GE, Henikoff S. 2012. Surveying the epigenomic landscape, one base at a time. Genome Biol 13: 250.

72. Zhang H, Emerson DJ, Gilgenast TG, Titus KR, Lan Y, Huang P, Zhang D, Wang H, Keller CA, Giardine B, et al. 2019. Chromatin structure dynamics during the mitosis-to-G1 phase transition. Nature 576: 158–162. 10.1038/s41586-019-1778-y.

73. Zhou CY, Dekker B, Liu Z, Cabrera H, Ryan J, Dekker J, Heald R. 2023. Mitotic chromosomes scale to nuclear-cytoplasmic ratio and cell size in Xenopus. Elife 12. https://elifesciences.org/articles/84360.

